# Ablation of tanycytes of the arcuate nucleus and median eminence increases visceral adiposity and decreases insulin sensitivity in male mice

**DOI:** 10.1101/637587

**Authors:** Sooyeon Yoo, David Cha, Soohyun Kim, Lizhi Jiang, Mobolanie Adebesin, Andrew Wolfe, Ryan Riddle, Susan Aja, Seth Blackshaw

## Abstract

Tanycytes are radial glial cells located in the mediobasal hypothalamus. Recent studies have proposed that tanycytes play an important role in hypothalamic control of energy homeostasis, although this has not been directly tested. Here, we report the phenotype of mice in which tanycytes of the arcuate nucleus and median eminence were conditionally ablated. Although the CSF-hypothalamic barrier was rendered more permeable, the blood-hypothalamic barrier was not altered. The metabolic effects of tanycyte ablation were likewise moderate. However, we consistently observed a significant increase in visceral fat distribution accompanying insulin insensitivity, but only in male mice, and without an effect on either body weight or food intake. A high-fat diet accelerated overall body weight gain in tanycyte-ablated mice, but the development of visceral adiposity and insulin insensitivity was attenuated. These results clarify the extent to which tanycytes regulate energy metabolism, and indicate a role for tanycytes in controlling body adiposity.

## Introduction

Tanycytes are radial glia that line the ventricular wall of the mediobasal hypothalamus. These are divided into alpha- and beta-subtypes, based on their dorsoventral position along the third ventricle. They extend elongated foot processes away from the ventricle, which terminate either on blood vessels in the hypothalamic parenchyma or at the ventral pial surface [1]. Beta-tanycytes reside outside the blood-brain barrier, and directly contact fenestrated capillaries in the median eminence [2]. Their location at the interface between the hypothalamus and circulatory system allows tanycytes to regulate a broad range of processes relevant to hypothalamic physiology. These include control of the blood-hypothalamus barrier [3,4], energy and nutrient sensing [5–7], and regulation of neurohormone release [8,9]. Tanycytes also express many markers of neural progenitor cells, and have been reported to show limited neurogenic potential [10–12].

Recently, there has been growing interest in the potential role of tanycytes in regulating energy metabolism [5,13,14]. Tanycytes have been reported to directly sense glucose and amino acid levels in CSF and blood [15–17], to actively transport leptin into the hypothalamus [18], and to regulate production and release of thyroid releasing hormone (TRH) [9,19]. However, these studies have not directly investigated whether tanycytes are indeed necessary for regulating these processes. It has thus far not been possible to selectively ablate tanycytes, and to directly observe the resulting effects on metabolism, or any other physiological process. This has been the major obstacle to an accurate understanding of their roles in normal hypothalamic physiology.

The development of transgenic mice that express tamoxifen-inducible tanycyte-specific *Cre* recombinase has made it possible to analyze the acute physiological effects of tanycyte ablation [20]. When crossed to mice expressing *Cre*-dependent *Eno2-lsl-DTA* [21], we observed a rapid, selective and essentially complete ablation of tanycytes of the arcuate nucleus and median eminence (ArcN-ME), which represent all beta-tanycytes, as well as a subset of alpha-2 tanycytes. We have used these mice to directly investigate the roles of tanycytes of the ArcN-ME in acute regulation of hypothalamic barrier function, leptin signaling, neurohormone release, fat distribution, insulin sensitivity and energy balance.

## Results

### 1. Conditional ablation of tanycytes in the ArcN-ME

To investigate the potential physiological contribution of tanycyte-derived neurogenesis [12,22–24], we generated *RaxCreER;Eno2-lsl-DTA* mice, with the intention of selectively ablating tanycyte-derived neurons without killing the tanycytes themselves. Analysis of *Eno*2 using fISH, however, revealed high levels of *Eno2* mRNA expression in hypothalamic neurons, along with unexpected low levels of expression in both beta- and ventral alpha-2 tanycytes (Fig. 1A). Expression of *Eno2* in tanycytes was subsequently confirmed using our scRNA-Seq data [25,26]. Because the generation and maturation of neurons in adults is a slow process [27], this makes it possible to use *RaxCreER;Eno2-lsl-DTA;Ai9* mice to study the acute physiological effects of tanycyte ablation.

**Figure 1.**
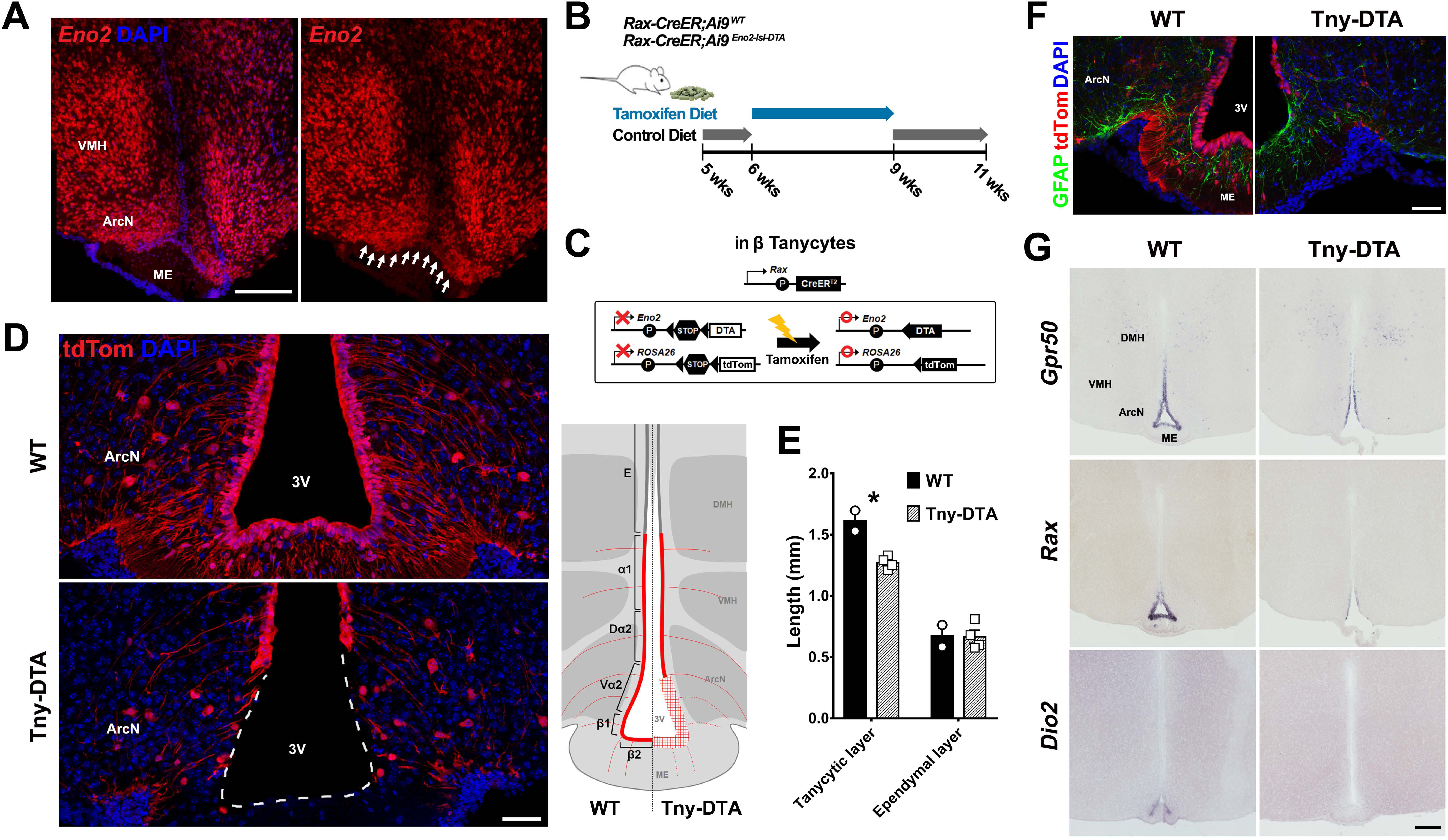
Tanycyte-specific ablation targeting the region of ArcN and ME. **(A)** *Eno2* mRNA expression in tanycytes located near the ArcN and ME (white arrows). **(B)** Schematic diagram showing the time schedule for administration of tamoxifen diet. Five-week old *RaxCreER;Ai9(WT) or RaxCreER;Eno2-lsl-DTA;Ai9(Tny-DTA)* mice were fed with control diet which, aside from lacking tamoxifen, has the same dietary components as the tamoxifen diet, for a week followed by 3 weeks of tamoxifen diet. The mice had at least one week of washout to allow for clearance of tamoxifen before experiments were conducted. **(C)** Schematic diagram showing the approach used to conditionally ablate tanycytes of the ArcN and ME. By using the *RaxCreER* line, *Cre* recombination induces *tdTom* expression in all subtypes of tanycyte while *DTA* expression is specific in *Eno2*-positive beta-tanycytes and ventral alpha-2 tanycytes. **(D)** Immunohistochemistry for tdTom showing nearly complete ablation of tanycytes in the ArcN and ME. **(E)** Quantification of reduction of tdTom-positive tanycytes following tamoxifen-induced ablation. **(F)** Immunohistochemistry for GFAP indicating reactive astrogliosis in the ventricular region following tanycyte ablation. **(G)** mRNA *in situ* hybridization using probes for tanycyte marker genes *Gpr50, Rax* and *Dio2*. Scale bar: 50 µm (D,F), 200 µm (A,G).

Loss of tanycytes in *RaxCreER;Eno2-lsl-DTA;Ai9* mice was observed at around 14-16 days after the start of dietary tamoxifen (Tm) administration (data not shown). The adult stem cell-like regenerative potential of tanycytes, along with this preliminary data, led us to use a 3 week dietary tamoxifen treatment (Fig. 1B & C). This was sufficient to lead to a near-complete ablation of tanycytes in ArcN-ME, as determined by loss of tdTom fluorescence (Fig. 1D). This corresponds to all of the beta-tanycytes and most of the ventral alpha-2 tanycytes, while the alpha-1 and dorsal alpha-2 tanycytes of the ventromedial and dorsomedial hypothalamus were spared, as were the ependymal cells of the dorsomedial hypothalamus (Fig. 1E). Reactive gliosis, as measured by increased level of GFAP immunostaining, was evident in the ventricular region of the ArcN-ME following tanycyte ablation (Fig. 1F).

Loss of tanycyte-specific markers such as *Rax, Gpr50* and *Dio2* was observed in the ArcN-ME (Fig. 1G). However, expression of hypothalamic neuropeptides in hypothalamic regions in direct contact with tanycytes, such as *Npy, Cart, Gal* and *Sst* (Fig. S1A). Likewise, the size of hypothalamic nuclei that are directly contacted by tanycytes was unaffected, with the exception of the ME, which showed a significant decrease in overall size (p<0.01) (Fig. S1B). However, axons of neurosecretory cells targeting the ME were not altered (Fig. S1C). Furthermore, expression of hypothalamic neurohormones such as *Trh, Oxt* and *Avp* was also unaffected by tanycyte ablation (Fig. S1D).

### 2. ArcN-ME tanycytes are required for the CSF-hypothalamus barrier but not the blood-hypothalamus barrier

We next investigated whether hypothalamic barrier function was affected by ablation of tanycytes in the ArcN-ME. Previous studies have suggested a role for beta-tanycytes in the formation of the blood-hypothalamus barrier [2,3] and control of its permeability in response to dietary signals [4]. Both alpha-2 and beta-tanycytes have also been implicated in formation of the CSF-hypothalamus barrier [28] and in active transport of small molecules and proteins between the CSF and hypothalamic parenchyma [15,17]. We examined the permeability of both the blood-hypothalamus and CSF-hypothalamus barriers in tanycyte-ablated mice using Evans Blue, delivered via both intravenous (i.v.) and intracerebroventricular (i.c.v) injections [3]. Tanycyte-ablated mice showed no difference in blood-hypothalamic barrier permeability, as measured by Evan Blue staining after i.v. delivery, with reduced absolute levels of ME staining which mainly reflected the overall reduction in ME volume that is seen in tanycyte-ablated mice (Fig. 2A & B). In contrast, i.c.v. delivery of Evans Blue demonstrated substantially increased permeability in ArcN of tanycyte-ablated mice, while the staining in the ME was still strong near the ventricular floor (Fig. 2C & D).

**Figure 2.**
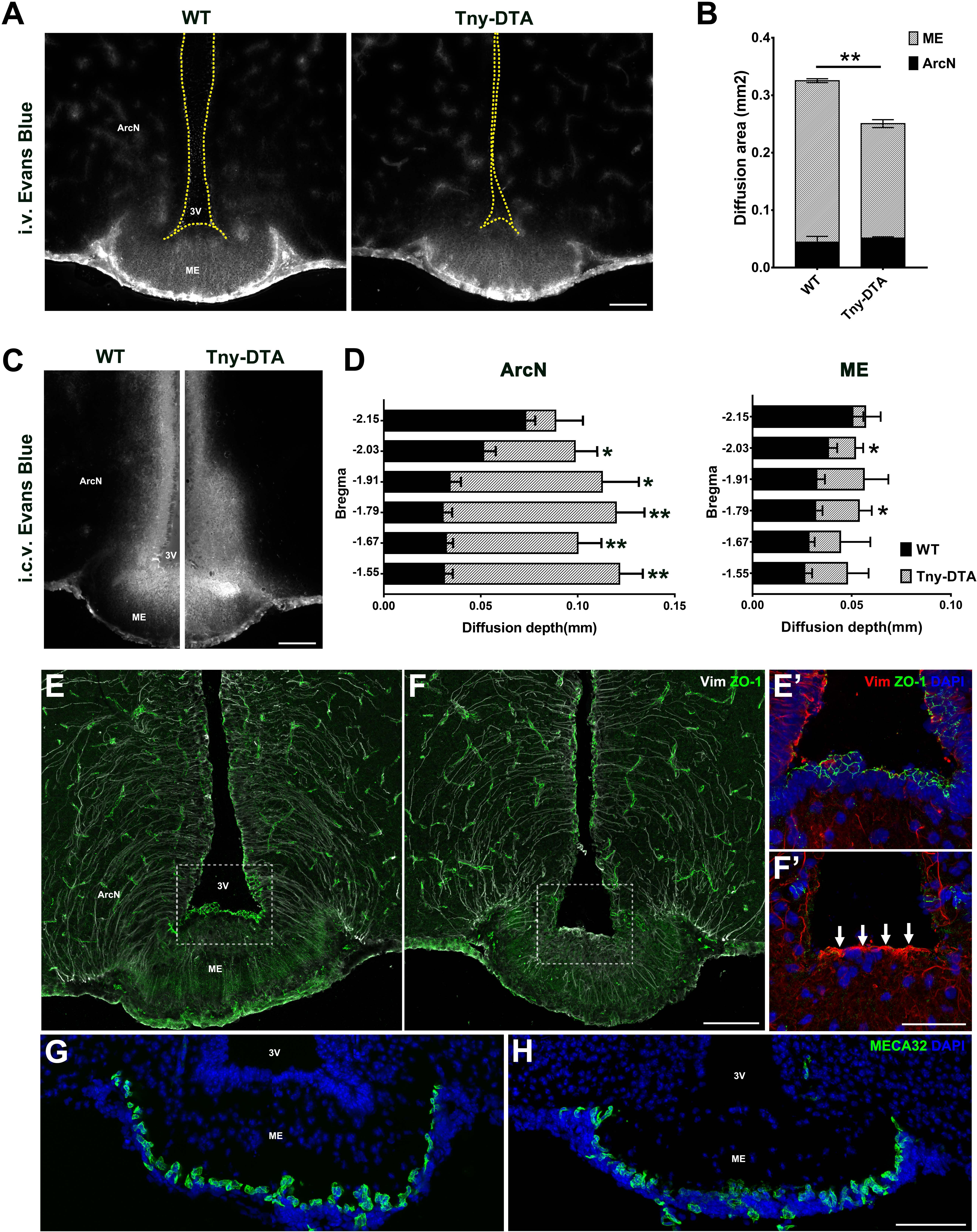
Characterization of a possible barrier defect in tanycyte-ablated mouse brain. **(A)** Representative images of Evans Blue staining in the ventromedial hypothalamus 20 min after i.v. injection. **(B)** Quantification of the area stained with Evans Blue, which is mostly restricted to the ArcN and ME. **(C)** Representative images of Evans Blue staining 5 min after i.c.v. injection. **(D)** Quantification of the penetration of Evans Blue staining into the ArcN or ME in the hypothalamic sections corresponding to the indicated bregma. **(E**,**F)** Immunohistochemistry for the tight junction protein ZO-1 (green) and intermediate filament Vimentin (Vim, white). The staining patterns for ZO-1 and Vim were lost following tanycyte ablation. Increased Vim expression (white arrows) in the third ventricular floor reveals a glial scar formation. **(E’**,**F’)** are the higher magnification images of the boxed area in (E,F). **(G**,**H)** Immunostaining against the fenestrated capillary marker, MECA32 (green). Scale bar: 100 µm (A,C,F,H), 50 µm (F’). \**p*<0.05, \*\**p*<0.005.

Immunostaining for the tight junction marker ZO-1, which is expressed in tanycytes and reported to maintain hypothalamic barrier function, is lost in both ArcN and ME tanycytes in *RaxCreER;Eno2-lsl-DTA* mice (Fig. 2E & F). In contrast, Vimentin immunoreactivity is increased in the ventricular zone of the ME (Fig. 2E’ & F’). This is also seen for GFAP immunostaining (Fig. 1F), indicating reactive astrogliosis. Immunostaining for MECA32, a marker of fenestrated capillaries, revealed no change in the number of immuno-positive cells in ME between the two groups (Fig. 2F). Together, these data demonstrate that, while tanycytes are necessary for maintenance of the CSF-hypothalamus barrier, they are not required to maintain the blood-hypothalamus barrier.

A previous report suggested that beta-2 tanycytes were involved in active transcytosis of leptin from blood to the hypothalamus [18]. Our recent study, however, has demonstrated that leptin receptors are not detectably expressed in tanycytes, and are not required for leptin transport into hypothalamus [26]. To further investigate whether tanycytes of the ArcN-ME might regulate hypothalamic leptin signaling, we performed intraperitoneal (i.p.) leptin injection in both control and tanycyte-ablated mice. We observed no differences in pSTAT3 staining in neurons of any hypothalamic region that were directly in contact with tanycytes (Fig. 3A & B). This data further confirms that tanycytes are not involved in active transport of leptin from blood to the hypothalamus.

**Figure 3.**
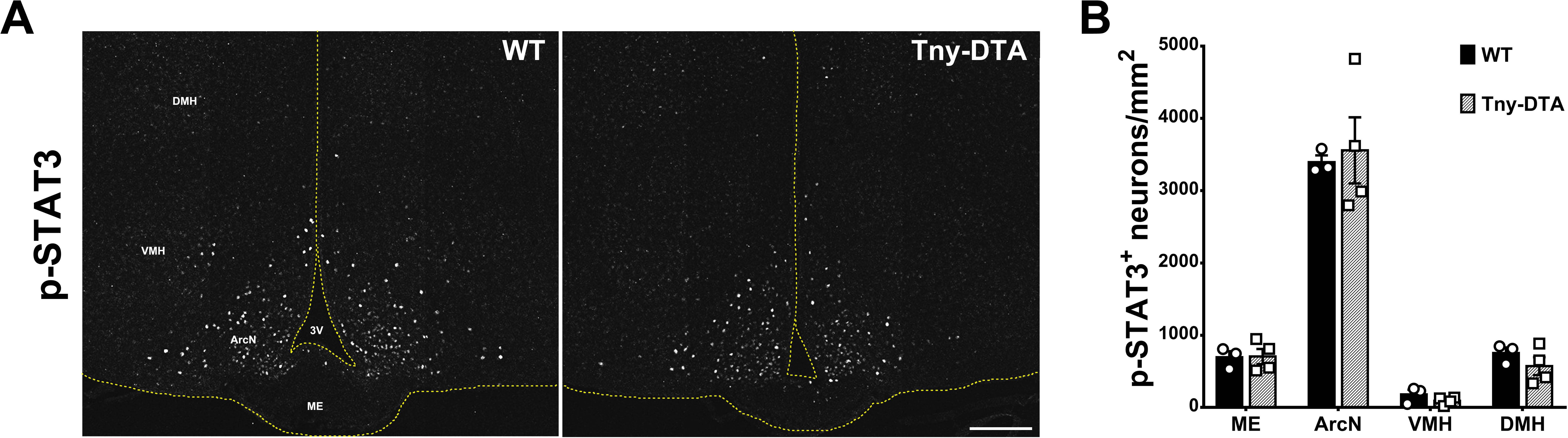
Leptin-induced STAT3 phosphorylation in the hypothalamus of control and tanycyte-ablated mice. **(A)** Representative images of pSTAT3 immunohistochemistry 15 min after i.p. leptin injection. **(B)** Quantification of the pSTAT3-positive neurons in the images shown (A), n= 3-4. DMH, dorsomedial nucleus; VMH, ventromedial nucleus; ArcN, arcuate nucleus; ME, median eminence; 3V, third ventricle. Scale bar: 100 µm (A).

### 3. Ablation of ArcN-ME tanycytes led to increased fat mass without body weight change, particularly in male mice

To more broadly investigate the contribution of tanycytes of the ArcN-ME to energy balance and other metabolic outcomes, we first analyzed food intake and body weight in control and tanycyte-ablated mice. We initially used a prolonged tamoxifen treatment to investigate the potential cumulative effects of loss of both tanycytes and tanycyte-derived cells, in both males and female mice. Both control and tanycyte-ablated mice were fed dietary tamoxifen for a period of over 50 days (Fig. 4A). While no difference in body weight was observed, a significant increase in fat mass was observed in male, but not in female mice, within 4 weeks following initiation of the tamoxifen diet (Fig. 4B-F). In particular, males had a substantial increase in perigonadal fat mass (Fig. 4G).

**Figure 4.**
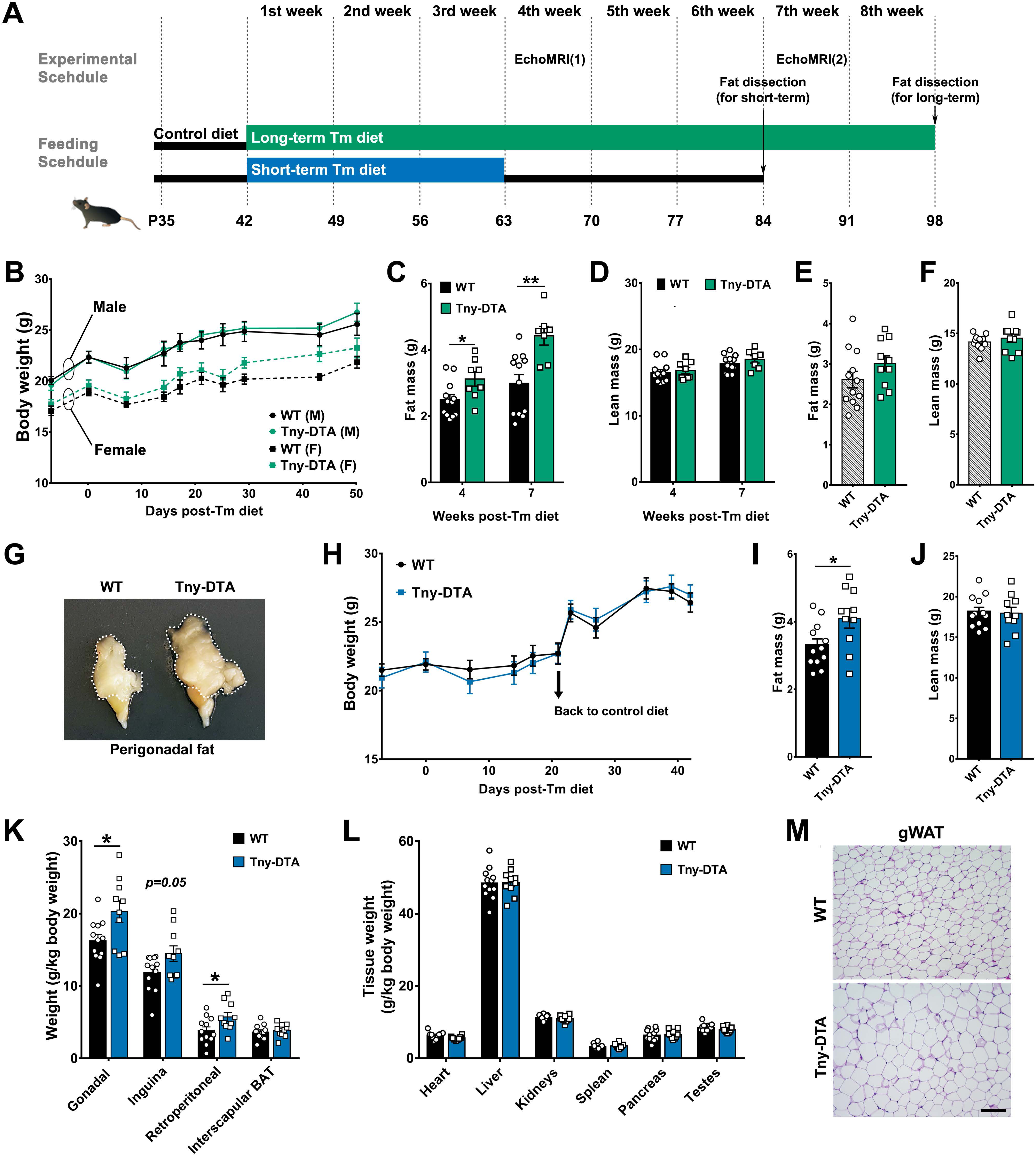
Effect of tanycyte-ablation on body weight and fat mass. **(A)** Strategy for the long-term and short-term dietary tamoxifen administration and experimental schedule. **(B)** Body weight change during the long-term dietary tamoxifen administration. **(C**,**D)** Fat and lean mass measured by EchoMRI at 4 weeks and 7 weeks following initiation of tamoxifen diet in males. **(E**,**F)** Fat and lean mass at 7 weeks following initiation of tamoxifen diet in females. **(G)** A representative image of perigonadal fat pad of wildtype (WT) and tanycyte-ablated mice (Tny-DTA). **(H)** Body weight change with the short-term tamoxifen feeding strategy. **(I, J)** Fat and lean mass measured by EchoMRI 4 weeks following initiation of tamoxifen diet. **(K, L)** Weight of dissected fat pads and other internal organs. **(M)** Representative H&E staining of gonadal white adipose tissue (WAT). Scale bar: 100um (M). \**p*<0.04, \*\**p*<0.01.

The increase in fat content in male mice seen within 4 weeks after initiating tamoxifen diet prompted us to investigate whether this effect was the result of acute tanycyte ablation, rather than cumulative effects of long-term ablation. Following 3 weeks of tamoxifen diet (as described in Fig.1B), we also did not observe differences in body weight during tamoxifen administration, or in the 2 weeks of return to control diet in male mice (Fig. 4H). However, a clear increase in body fat content was observed, while lean body mass was not affected (Fig. 4I & J). Increased perigonadal and retroperitoneal fat depots were observed, with a similar trend for inguinal fat (p=0.05) (Fig. 4K). Perigonadal adipocytes were larger in tanycyte-ablated mice than in wildtype mice (Fig. 4M). We did not find significant differences in brown adipose tissue (Fig. 4K) or in the sizes of any other internal organs between the two mouse groups (Fig. 4L).

### 4. Increased fasting-induced food intake and reduced insulin sensitivity is observed in ArcN-ME tanycyte-ablated male mice

In order to explore responses to states of negative and positive energy balance, we measured food intakes in control and tanycyte-ablated animals fed *ad libitum*, and then re-fed after an overnight food deprivation, during the long-term ablation (Fig.5A). In male mice maintained on tamoxifen for 4 weeks, no differences in food intake were observed in mice fed *ad libitum*, or following refeeding after the fast (Fig. 5B), indicating that food intake was not altered in the early phase of fat accumulation. However, after 8 weeks of tamoxifen treatment, male tanycyte-ablated mice showed hyperphagia, particularly in response to food deprivation (Fig. 5C).

**Figure 5.**
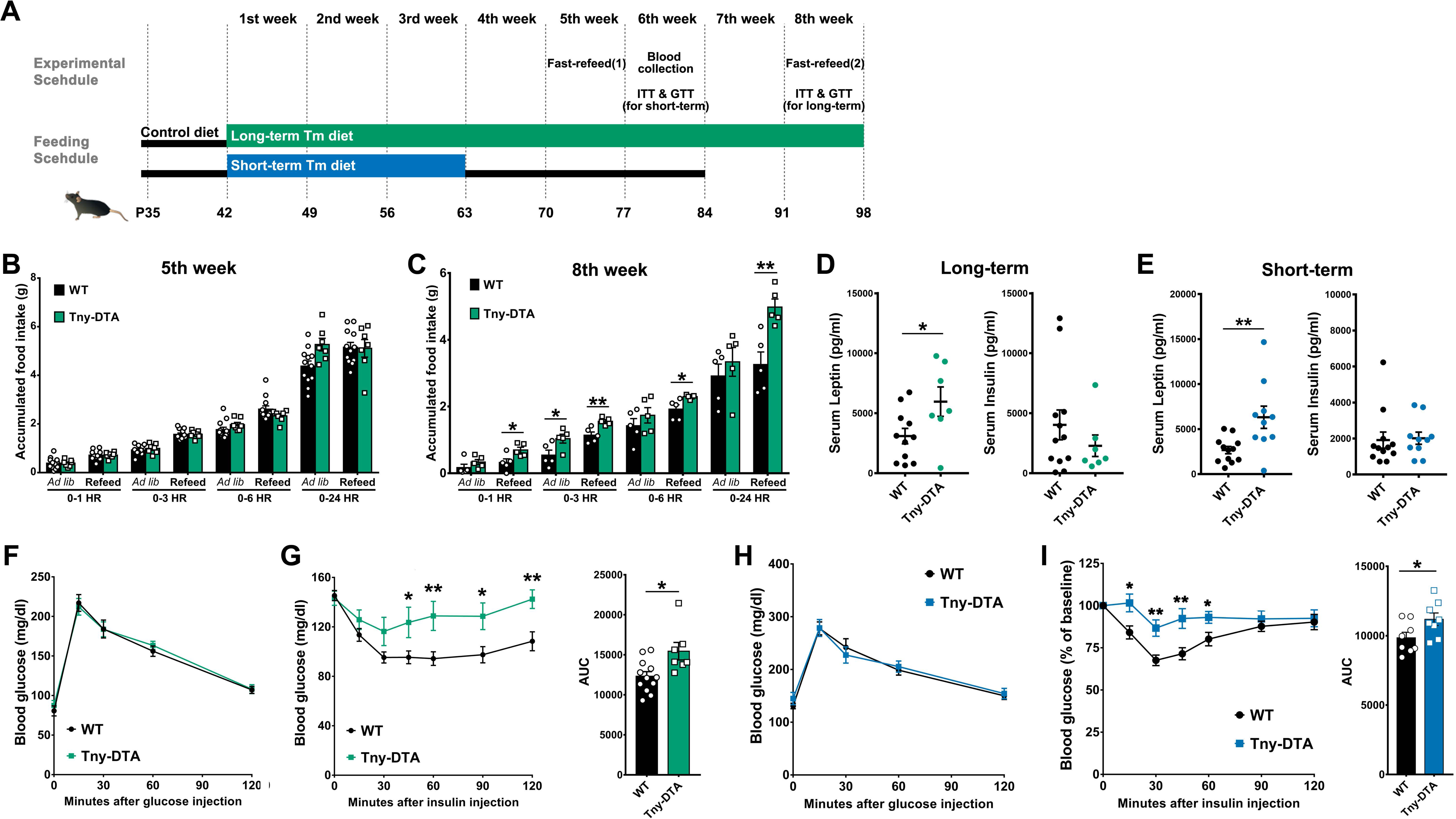
Effects of tanycyte-ablation on food intake, metabolic hormone level and glucose/insulin sensitivity. **(A)** Experimental schedule during the dietary tamoxifen administration. **(B**,**C)** *Ad libitum* (*Ad lib*) or 18h fasting-induced (re-feed) food intake at 5 weeks (B) and 8 weeks (C) following initiation of tamoxifen diet. **(D, E)** Serum leptin and insulin level measured by Luminex assay using a metabolic bead panel. **(F)** Glucose tolerance test in long-term tamoxifen fed mice and area under the curve (AUC). **(G)** Insulin tolerance test in long-term tamoxifen-fed mice. **(H)** Glucose tolerance test in short-term tamoxifen-fed mice. **(I)** Insulin tolerance test in short-term tamoxifen fed mice and area under the curve (AUC). \**p*<0.04, \*\**p*<0.01.

To further investigate mechanisms that might contribute to the hyperphagia, we analyzed blood hormone levels in male mice subjected to both short- and long-term tamoxifen treatments. Mice that underwent either short-term or long-term tamoxifen treatment showed significant increase in serum leptin, consistent with the increased fat contents, but showed no change in serum insulin (Fig. 5D & E). Satiety hormones from gut, pancreatic polypeptide (PP) and peptide YY (PYY), were increased significantly in tanycyte-ablated mice following long-term tamoxifen treatment (Fig. S3A & B). No changes were observed in serum levels of leptin and insulin in females (Fig. S2C).

We did not observe effects on glucose tolerance after tanycyte ablation induced by short-term or long-term tamoxifen treatments (Fig. 5F & H). However, tanycyte-ablated male mice exhibited substantially reduced sensitivity to insulin under both conditions (Fig. 5G & I), consistent with the increased adiposity in these animals. In contrast, females exhibited no differences in food intake, glucose tolerance or insulin sensitivity (Fig. S2A,B,D & E).

To further investigate the mechanisms mediating the acute increase in fat contents, and to test the more general role of tanycytes in regulation of release of hypothalamic and pituitary hormones, we expanded our analysis to test serum levels of a broad panel of factors (Fig. S3, S4 & S5). Surprisingly, few consistent changes in hormone levels were observed. Significant increases in serum TSH and neurotensin levels were observed in tanycyte-ablated mice following short-term treatment (Fig.S4J & O), although neither hormone was affected in tanycyte-ablated mice following long-term tamoxifen treatment (Fig. S3J & O). Because tanycytes produce thyroid hormone converting enzyme [29], and have been linked to regulation of TRH release from hypothalamic neurosecretory cells [9,19,30], we further investigated whether short-term tanycyte-ablation compromised the response of the hypothalamic-pituitary-thyroid axis to an acute cold challenge. While serum TSH levels were consistently elevated following 2 hour cold exposure, no significant difference was observed between control and tanycyte-ablated mice (Fig. S6).

Decreased serum oxytocin levels were observed in tanycyte-ablated mice following long-term tamoxifen treatment (Fig. S3M). No changes were observed in serum hormone levels in females (Fig.S5), with the surprising exception of growth hormone (GH), which showed two-fold higher levels following tanycyte-ablation (Fig. S5J). Since females do not show an overall increase in body weight or lean mass, the significance of this observation is unclear.

To further investigate the mechanism underlying the adiposity observed in tanycyte-ablated mice, we conducted indirect calorimetry studies on male mice that had undergone short-term tanycyte ablation (Fig.S7A). As observed previously, there were no differences in body weight or caloric intake between controls and tanycyte-ablated mice (Fig. S7B & C). Likewise, there were no differences in VO_2_ (or VCO2, data not shown) or rates of energy expenditure (Fig. S7E & F). However, a close examination of respiratory exchange ratio (RER) suggested a subtle shift away from fat oxidation, during the later portion of the dark phase and during the light phase, when compared to control mice (Fig. S7D).

### 5. Effects of high-fat diet on tanycyte-deficient male mice

The increased fat contents and decreased insulin sensitivity in tanycyte-deficient mice that had been fed a low-fat diet raised the possibility that these effects might be enhanced on an obesogenic high-fat diet (HFD). To address this, we placed both control and tanycyte-ablated male mice on a high-fat diet immediately following 3 weeks of feeding with tamoxifen-supplemented low-fat diet (Fig. 6A). Tanycyte-ablated mice showed increased body weight gain on HFD compared with control mice (Fig. 6B). HFD produced a robust increase in overall fat mass in both groups, but tanycyte-ablated mice showed statistically greater overall fat mass than wildtype mice (Fig. 6C). However, none of the individual depots of white adipose tissue showed increased mass in tanycyte-ablated mice compared with controls (Fig. 6D), in contrast to the substantial increases seen in ablated mice maintained on regular low-fat tamoxifen diet (Fig. 4K).

**Figure 6.**
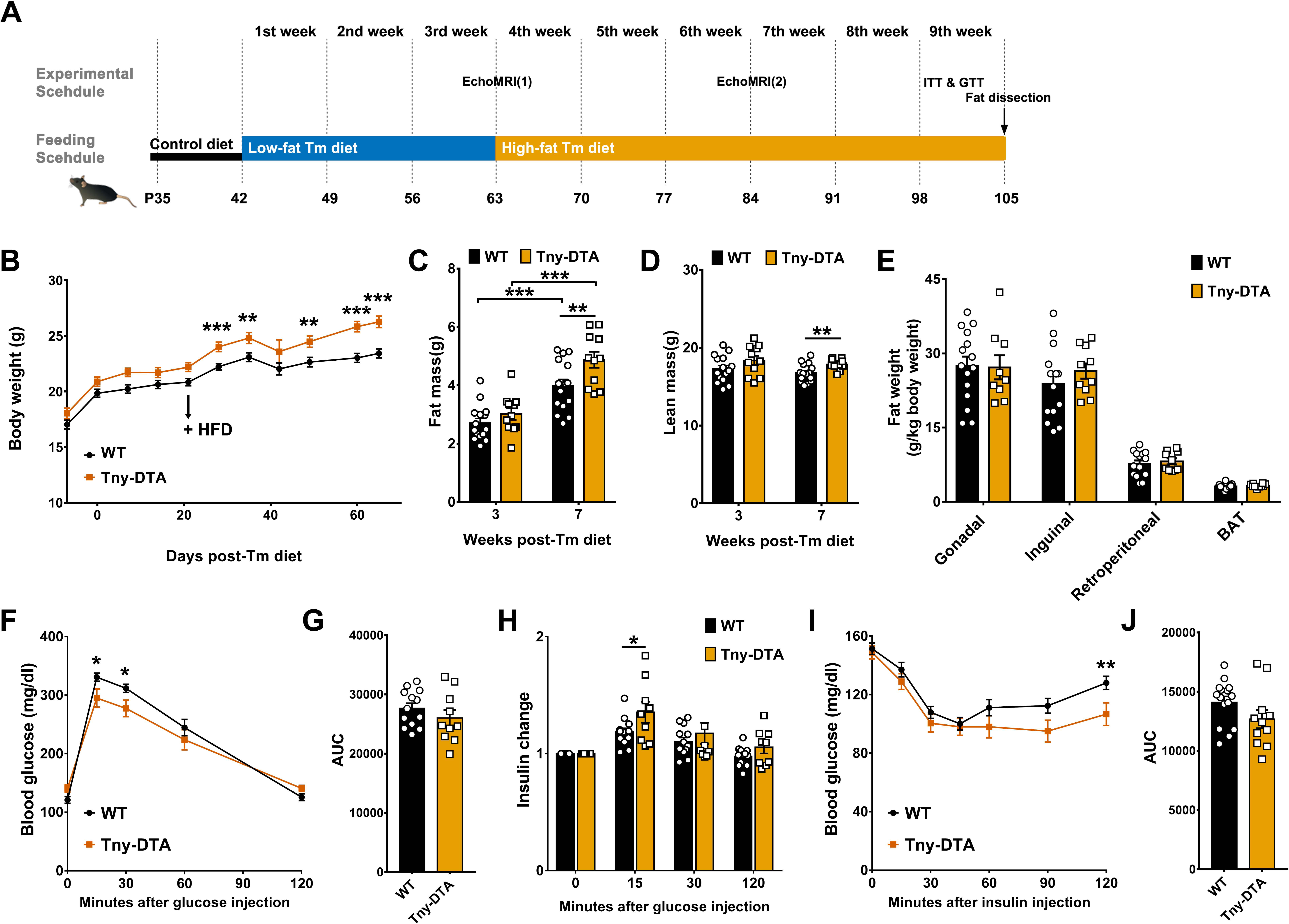
Effect of high-fat diet on metabolic phenotype in the tanycyte-deficient mice. **(A)** Experimental schedule during the high-fat tamoxifen diet administration. **(B)** Body weight change during the tamoxifen administration. **(C**,**D)** Fat and lean mass measured by EchoMRI before and after 3 weeks of high-fat tamoxifen diet **(E)** Weight of dissected fat pads. **(F)** Glucose tolerance test in high-fat tamoxifen diet fed mice. **(G)** Area under the curve (AUC) quantification of (F). **(H)** Serum insulin level change during the course of the glucose tolerance test in (F). **(I)** Insulin tolerance test in mice fed with high-fat tamoxifen diet. **(J)** Area under the curve (AUC) quantification of (I). \**p*<0.05, \*\**p*<0.02, \*\*\**p*<0.002.

Unlike ablated mice maintained only on low-fat tamoxifen diet (Fig. 5H), HFD-fed tanycyte-deficient mice showed slight but significant improvements in glucose tolerance at early time points (Fig. 6F & G), consistent with increased serum insulin 15 min after glucose injection (Fig. 6H). Tanycyte-ablated male mice showed prolonged effects of insulin action in an insulin tolerance test on HFD (Fig. 6I & J). This effect was also more moderate than that observed on regular tamoxifen diet which showed decreased insulin sensitivity at earlier time points than mice on high-fat tamoxifen diet. (Fig. 5I).

## Discussion

In this study, we have investigated the cellular and physiological effects of selective ablation of tanycytes of the ArcN and ME. Although tanycytes in these hypothalamic regions have been proposed to control a broad range of physiological processes, including blood-hypothalamic barrier formation, control of hypothalamic leptin signaling and regulation of the neuroendocrine axis, our findings revealed surprisingly subtle phenotypes in tanycyte-ablated mice, which is primarily restricted to a male-specific increase in adiposity and a decreased insulin sensitivity, without body weight change.

Perhaps most surprisingly, complete ablation of beta-tanycytes did not lead to an increased permeability of the blood-hypothalamus barrier, although it led to a clear increase of Evans Blue permeability from the CSF into the ArcN parenchyma, along with minor leakage into the ME. This is consistent with a previous report that tanycytes regulate CSF-hypothalamic barrier function [28]. This likely implies a central role for other cell types, such as endothelial cells and pericytes, in regulating permeability between the ArcN and ME. The finding that hypothalamic leptin signaling was unaffected by absence of beta-tanycytes is consistent with our recent findings that LepR expression in tanycytes is not necessary for leptin transport from blood to hypothalamic parenchyma [26].

We likewise observed only modest and transient effects on the neuroendocrine axis. beta-tanycyte have been implicated in active regulation of both GnRH and TRH synthesis and/or release [8,9,30]. Tanycyte foot processes have been reported to physically inhibit GnRH release in female, but to retract during estrus to allow GnRH release into the portal circulation [8,31]. While we did not observe significant changes in serum levels of FSH or LH in either sex, these results are difficult to interpret due to disruptive effects of prolonged tamoxifen exposure on the hypothalamic-pituitary-gonadal axis.

Beta-tanycytes strongly express *Dio2*, an enzyme that converts T4 to T3, which is known to negatively regulate transcription of *Trh* [32]. Furthermore, beta-tanycytes release exoenzymes which degrade TRH [19,30]. One might therefore predict that ablation of beta-tanycytes would lead to a global activation of the hypothalamic-pituitary-thyroid (HPT) axis [33]. We did not, however, observe any obvious changes in hypothalamic *Trh* expression in tanycyte-ablated mice (Fig. S1). While we observed an increase in serum TSH in male mice subjected to short-term tanycyte ablation (Fig. S4), consistent with HPT hyperactivation, no significant change in serum TSH was observed in either male mice subjected to longer-term tanycyte ablation or in female mice (Fig. S3 & 5). Furthermore, we did not observe statistically significant differences in elevated serum levels of TSH following acute cold exposure between the control and tanycyte-ablated mice (Fig. S6). This suggests that any role beta-tanycytes play in regulating the HPT is relatively modest and also sex-specific, and that disruption of beta-tanycyte function is rapidly compensated for by changes in other tissues.

Beta-tanycytes have been reported to sense glucose and amino acids [15–17], and have been proposed to act as essential central integrators of metabolic regulation [5,13,14]. Nevertheless, tanycyte ablation did not result in dramatic changes in regulation of energy balance in normal and obesogenic dietary conditions. Tanycyte-ablated mice did not show changes in overall body weight or food intake in the 4 weeks following tanycyte ablation. However, we observed a significant increase in adiposity, that is most prominent in visceral fat, in male mice at this stage (Fig. 4G & K). Within 4 weeks following tanycyte ablation, decreased insulin sensitivity was observed, along with increased serum leptin levels. However, glucose tolerance was unaffected, suggesting that these changes may be secondary to the increased adiposity. In females, however, no significant changes in adiposity, serum leptin and insulin sensitivity were observed (Fig. 4E & Fig. S2). The molecular mechanisms underlying these acute metabolic changes and their sexual dimorphism are yet unclear. However, the data suggest that there is a direct or indirect involvement of tanycytes in control of fat utilization, as shown by the slightly higher RER overall during typically fat-oxidizing phases of the photoperiod in tanycyte-deficient male mice (Fig. S7D).

Increased adiposity in males became more prominent at 7 weeks following tanycyte ablation, and yet still without overall body weight change (Fig. 4B & C). This was accompanied by an exacerbated hyperphagic response to prolonged fasting, although *ad libitum* food intake was unaffected (Fig. 5C). This enhanced re-feeding implies that longer-term tanycyte ablation disrupts both normal central integration and the response to signals of negative energy balance. This could result from a variety of mechanisms, ranging from the absence of sensation of small molecules by tanycytes to the disruption of generation of tanycyte-derived neurons or glia [11,17].

In conclusion, tanycytes of the ArcN-ME have a specific role in fat metabolism rather than food intake and body weight control which would be more closely related to the physiological functions of tanycytes recently reported [7,9,13]. This does not rule out a potentially important role for tanycytes in influencing these processes, but demonstrates that other cells can compensate for the loss of tanycyte function. It also remains possible that the residual tanycytes, which would be alpha-1 and a subset of alpha-2 subtypes, could be major players in the expected metabolic phenotypes [34,35] Furthermore, we have only tested a limited number of potential environmental conditions, and a more prominent role for tanycytes may be revealed in response to other physiological challenges. Finally, there are many other physiological processes that may be tanycyte-regulated that were not tested in this study. Notably, tanycytes have been reported to play a critical role in biological timing and reproductive regulation in species that show strong photoperiodism [36–38], and it is possible that related behaviors may be under tanycyte-dependent regulation in mice.

## Acknowledgements

We thank Dr. Shigeyoshi Itohara for generously providing the *Eno2-lsl-DTA* mice. We thank W. Yap, J. Ling, T. Hoang, and C. Santiago for their comments on the manuscript. This work was supported by grant DK108230 to S.B. and a Maryland Stem Cell Postdoctoral Research Fellowship to S.Y.

## Materials and Methods

### Animals

*Rax-CreERT2* mice gene were generated in the laboratory were crossed with Ai9 (*R26-CAG-lsl-tdTom*, JAX #007909) [20] (JAX#025521), and then bred with *Eno2-lsl-DTA* mice [39] to induced the tanycyte-specific ablation. To induce Cre recombination, the mice were fed with commercial tamoxifen-containing diet (250 mg/kg diet, Envigo Teklad diets #TD. 130856) starting at P42 for the indicated periods of time to produce and maintain the tanycyte ablation. To aid acclimation from regular diet presented at weaning (TD.2018, 3.1 kcal/g; 24 kcal% protein, 18 kcal% fat, 58 kcal% carbohydrate) to the tamoxifen-containing diets, mice were fed from P35 until P42 on a diet that was the same energy density, but with macronutrients otherwise matched to the subsequent tamoxifen-containing diet (control diet without tamoxifen: 3.1 kcal/g; 19.7 kcal% protein, 70.6 kcal% carbohydrate, 9.7 kcal% fat)(Envigo Teklad diet #TD. 07570). For the short-term ablation study, the mice were switched back to this control diet after 3 weeks, for the physiological phenotyping. For the high-fat diet feeding experiments, customized tamoxifen diets were based on either a very-high-fat diet (Research Diets, 5.2 kcal/g; 20 kcal% protein, 20 kcal% carbohydrate, 60 kcal% fat, supplemented tamoxifen at 403.1 mg/kg) or on a low-fat diet (Research Diets, 3.8 kcal/g; 20 kcal% protein, 70 kcal% carbohydrate, 10 kcal% fat supplemented with 248 mg/kg diet). Mice were initially housed in a specific pathogen free facility on a 14 h-10 h light/dark cycle (07:00 lights on – 19:00 lights off). One week prior to physiological studies, mice were transferred to the rodent metabolism core satellite facility at the Center for Metabolism and Obesity Research to acclimate to its vivarium on 12 h - 12 h light dark cycle (06:00 lights on - 18:00 lights off climates at both facilities were controlled within temperature and humidity ranges consistent with the Guide for the Care and Use of Laboratory Animals (8th ed.). All studies and procedures were performed under a pre-approved protocol by the Institutional Animal Care and Use Committee (IACUC) of the Johns Hopkins University School of Medicine.

### Tissue processing and Immunohistochemistry

Mice were anesthetized with intraperitoneal (i.p.) injection of Tribromoethanol/Avertin before transcardial perfusion with 1X PBS followed by 2% paraformaldehyde in 1X PBS. Brains were dissected and post-fixed in the same fixative for overnight at 4°C. After brief washing with 1X PBS, brains were incubated in 30% sucrose until brains sunk, then transferred and incubated in a 50:50 solution of 30% sucrose: O.C.T. embedding compound before being frozen. Brains were coronally sectioned at 25 µm thickness using a cryostat (Leica Biosystems) and stored in antifreeze solution at -20°C.

Sections were mounted on the Superfrost Plus slides (Fisher Scientific) before immunohistochemistry was conducted, and dried at room temperature for 30 min. For pSTAT3 staining, sections were sequentially incubated with 0.5% NaOH + 0.5% H_2_O_2_ for 20 min, 0.3% glycine for 10 min, and 0.03% SDS for 10 min, and blocked in 4% sheep serum/ 1% BSA/ 0.4% Triton X-100 in PBS for 1 hour at room temperature. Otherwise, sections were incubated with sodium citrate buffer (10mM Sodium citrate, pH 6.0) at 80°C for 30 min for antigen retrieval. After cooling for another 30 min at room temperature, sections were blocked in 10% sheep serum/ 0.1% Triton X-100 in PBS for 1 hour at room temperature. The sections imaged for Evans Blue staining were immediately treated with blocking solution before primary antibody incubation.

Antibodies used were as follows: rabbit anti-pSTAT3 (1:1000, #9145, Cell Signaling Technology), mouse anti-HuC/D (1:200, #A-21271 Invitrogen), rabbit anti-GFAP (1:500, #Z0334 DAKO), rabbit anti-ZO-1 (1:500, #61-7300 Invitrogen), rat anti-MECA32 (1: 200, DSHB), chicken anti-Vimentin (1:1000, #AB5733 Millipore), rat anti-red fluorescent protein (1:500, #5f8-100 Chromo Tek), rabbit anti-DsRed (1:500, #632496 Clontech), mouse anti-Neurofilament-M (1:50, #2H3 DSHB), donkey anti-rabbit Alexa Fluor^®^ 647 (1:500, #711-605-152, Jackson ImmunoResearch), goat anti-mouse IgG, Fcγ subclass 2b specific Alexa Fluor^®^ 488 (1:500, #115-545-207, Jackson ImmunoResearch), donkey anti-rabbit Alexa Fluor^®^ 488 (1:500, #711-545-152, Jackson ImmunoResearch), goat anti-chicken Alexa Fluor^®^ 647 (1:500, #A-21449 invitrogen), goat anti-rabbit Alexa Fluor^®^ 555 (1:500, #A-21428

Invitrogen), donkey anti-rat Alexa Fluor^®^ 488 (1:500, #712-545-153 Jackson ImmunoResearch).

After counterstaining with DAPI, sections were coverslipped using Vectashield antifade mounting medium (# H-1200, Vector Laboratories) before imaging on a Zeiss LSM 700 Confocal at the Microscope Facility (Johns Hopkins University School of Medicine).

### *In situ* hybridization

Chromogenic *in situ* hybridization was performed on fresh-frozen brain tissue as previously described [40,41]. Briefly, coronal 25 µm sections were prepared using a cryostat and fixed with 4% paraformaldehyde. After acetylation with triethanolamine (0.1M, pH 8.0) and acetic anhydride mixture (0.27%), sections were hybridized with digoxigenin-labeled probes at 70°C overnight. Unbound probes were washed out and sections were blocked with sheep serum followed by incubation with anti-digoxigenin antibodies conjugated to alkaline phosphatase (1:5000) overnight at 4°C. The combination of nitro-blue tetrazolium (NBT) and 5-bromo, 4-chloro, 3-indolylphosphate (BCIP) was used as chromogenic substrates of alkaline phosphatase. Color development was continued in the same amount of time between wildtype and tanycyte-ablated sections.

### Evans Blue injection

Mice given 2-3 weeks of reversion to matched control diet, after the 3 weeks of tamoxifen diet, were injected with sterile 1% Evans Blue dye in 0.9% saline through either intravenous (i.v.; 50µl) or intracerebroventricular (i.c.v.; 2µl) [4]. For i.v. injections, mice were decapitated 20 min after tail vein injections, and brains was dissected and immediately frozen in O.C.T. embedding compound. For i.c.v. injections, Evans Blue is injected into the lateral ventricle (y: - 0.3 mm, x: - 1mm, z: 2.5 mm) of anaesthetized mice (100 mg/kg ketamine, 10 mg/kg xylazine and 3 mg/kg acepromazine) using an infusion pump with 10-ul Hamilton syringe for 1 min. Infusion was confirmed by Evans Blue dye diffusion into the opened cisterna magna during the injection and the needle was removed 1 min after finishing the injection. The snap-frozen brains were coronally sectioned at 25 µm thickness and directly imaged under a fluorescent microscope (Keyence BZ-X700). The sections were then fixed in methanol/acetone (v/v) at -20°C for 1 min and further used for the immunohistochemistry as described above.

### Leptin injection

Leptin injection was performed as previously described [26]. Briefly, 3 mg/kg leptin was injected i.p., and 15 min later transcardial perfusion was performed with 1X PBS followed by 2% paraformaldehyde in 1X PBS. Previous studies of leptin-induced hypothalamic STAT3 phosphorylation studies [26,42,43] led us conclude that, if there is any leakage through the opened CSF-hypothalamus or blood-hypothalamus barrier, this would be evident within 15 min following leptin injection.

### Fasting-refeeding food intake study

Tamoxifen-fed male and female mice were individually housed and initially provided *ad libitum* access to water and food, in a vivarium on a 12 h-12 h light/dark cycle. A wire mesh floor was inserted into the home cage to collect spilled food onto a cage paper beneath the mesh. For the fasting-refeeding experiment, both groups of animals were fasted 18h (beginning halfway through the dark phase), and then refed for 24h (beginning at the start of the next dark phase). Food intake (corrected for dry, feces-free spillage) was measured at 1, 3, 6, and 24 hour time points during the course of refeeding. The *ad libitum* food intake data was obtained one day prior to the fast-refeed study with the same procedure used for food intake measurement.

### Luminex assay and ELISA

Mouse serum samples were collected from tail vein at between 14:00h and 16:00h. After brief exposure to heat to promote blood vessel dilation, the collection site was cleaned with 70% alcohol, and the mouse is restrained. Blood was collected using a sterile surgical blade (#12-460-440, Fisher Scientific) from the lateral tail vein. Blood flow was stopped by applying pressure with sterile gauze to achieve hemostasis. Metabolic hormone, pituitary hormone, and neuropeptide serum levels were measured by Luminex assay according to the manufacturer’s instructions (#MMHMAG-44K, #MPTMAG-49K, and #RMNPMAG-83K from Millipore Sigma). Serum insulin and testosterone levels were measured using ELISA kit purchased from Millipore (#EZRMI-13K) and Crystal Chem (#80552), respectively. For the acute cold exposure study, mice were transferred to a room maintained at 4°C for 2 hours at between 14:00 h and 16:00h. Blood was briefly collected immediately after cold exposure from the submandibular vein. Serum TSH levels were measured using ELISA kit purchased from Cloud-Clone Corp. (#CEA463Mu).

### Body composition analysis, Indirect calorimetry

Fat and lean masses were determined using a whole-body NMR instrument (EchoMRI, Waco, TX) at the Phenotyping Core of the Johns Hopkins University School of Medicine. Indirect calorimetry was conducted using a Comprehensive Lab Animal Monitoring System (Columbus Instruments) in the rodent metabolism core facility at the Center for Metabolism and Obesity Research of the Johns Hopkins University. Mice were acclimated and monitored in the individual test chambers for days, with a fourth day for data reporting.. Daily body weights and food intakes (corrected for spillage) were measured. Oxygen consumption (VO_2_, mL/kg/h) and carbon dioxide production (VCO_2_, mL/kg/h) were measured for each chamber every 10 min. Oxymax software (v. 4.02) calculated RER (VCO_2_/VO_2_) and energy expenditure (EE = CV x VO_2_, CV = 3.815 + (1.232 x RER)..

### Glucose and insulin tolerance tests (GTT, ITT)

For the glucose tolerance test, animals were fasted for 12 hours, and 0.1g/kg glucose was injected i.p.. Blood glucose levels during the test were measured using the tail-nick method and a glucometer (Novamax).In addition, up to 25 ul of blood was collected to obtain serum for insulin measurements with an ELISA kit. For the insulin tolerance test, food was removed 2 hour prior to 0.8 units/kg body weight insulin injection, and blood glucose was measured with a glucometer.

### Cell counting and statistical analysis

All pSTAT3 and HuC/D-double positive cell counts was performed blinded and blindly and manually, as performed described in our previous study [26]. The size (mm) of each hypothalamic nucleus and the length of tanycytic layer or Evans Blue penetration depth were measured using Image-J. All values are expressed as means ± S.E.M. Comparison was analyzed by unpaired, two-tailed Student’s *t*-test, and *p< 0.05* was considered statistically significant.

## References

1. Rodríguez E, Guerra M, Peruzzo B, Blázquez JL. Tanycytes: A rich morphological history to underpin future molecular and physiological investigations. J Neuroendocrinol. 2019;31: e12690.

2. Rodríguez EM, Blázquez JL, Guerra M. The design of barriers in the hypothalamus allows the median eminence and the arcuate nucleus to enjoy private milieus: the former opens to the portal blood and the latter to the cerebrospinal fluid. Peptides. 2010;31: 757–776.

3. Mullier A, Bouret SG, Prevot V, Dehouck B. Differential distribution of tight junction proteins suggests a role for tanycytes in blood-hypothalamus barrier regulation in the adult mouse brain. J Comp Neurol. 2010;518: 943–962.

4. Langlet F, Levin BE, Luquet S, Mazzone M, Messina A, Dunn-Meynell AA, et al. Tanycytic VEGF-A boosts blood-hypothalamus barrier plasticity and access of metabolic signals to the arcuate nucleus in response to fasting. Cell Metab. 2013;17: 607–617.

5. Ebling FJP, Lewis JE. Tanycytes and hypothalamic control of energy metabolism. Glia. 2018;66: 1176–1184.

6. Elizondo-Vega R, Cortes-Campos C, Barahona MJ, Oyarce KA, Carril CA, García-Robles MA. The role of tanycytes in hypothalamic glucosensing. J Cell Mol Med. 2015;19: 1471–1482.

7. Elizondo-Vega RJ, Recabal A, Oyarce K. Nutrient Sensing by Hypothalamic Tanycytes. Front Endocrinol. 2019;10: 244.

8. Parkash J, Messina A, Langlet F, Cimino I, Loyens A, Mazur D, et al. Semaphorin7A regulates neuroglial plasticity in the adult hypothalamic median eminence. Nat Commun. 2015;6: 6385.

9. Müller-Fielitz H, Stahr M, Bernau M, Richter M, Abele S, Krajka V, et al. Tanycytes control the hormonal output of the hypothalamic-pituitary-thyroid axis. Nat Commun. 2017;8: 484.

10. Haan N, Goodman T, Najdi-Samiei A, Stratford CM, Rice R, El Agha E, et al. Fgf10-expressing tanycytes add new neurons to the appetite/energy-balance regulating centers of the postnatal and adult hypothalamus. J Neurosci. 2013;33: 6170–6180.

11. Yoo S, Blackshaw S. Regulation and function of neurogenesis in the adult mammalian hypothalamus. Prog Neurobiol. 2018;170: 53–66.

12. Lee DA, Bedont JL, Pak T, Wang H, Song J, Miranda-Angulo A, et al. Tanycytes of the hypothalamic median eminence form a diet-responsive neurogenic niche. Nat Neurosci. 2012;15: 700–702.

13. Prevot V, Dehouck B, Sharif A, Ciofi P, Giacobini P, Clasadonte J. The Versatile Tanycyte: A Hypothalamic Integrator of Reproduction and Energy Metabolism. Endocr Rev. 2018;39: 333–368.

14. García-Cáceres C, Balland E, Prevot V, Luquet S, Woods SC, Koch M, et al. Role of astrocytes, microglia, and tanycytes in brain control of systemic metabolism. Nat Neurosci. 2019;22: 7–14.

15. Frayling C, Britton R, Dale N. ATP-mediated glucosensing by hypothalamic tanycytes. J Physiol. 2011;589: 2275–2286.

16. Benford H, Bolborea M, Pollatzek E, Lossow K, Hermans-Borgmeyer I, Liu B, et al. A sweet taste receptor-dependent mechanism of glucosensing in hypothalamic tanycytes. Glia. 2017;65: 773–789.

17. Lazutkaite G, Soldà A, Lossow K, Meyerhof W, Dale N. Amino acid sensing in hypothalamic tanycytes via umami taste receptors. Mol Metab. 2017;6: 1480–1492.

18. Balland E, Dam J, Langlet F, Caron E, Steculorum S, Messina A, et al. Hypothalamic tanycytes are an ERK-gated conduit for leptin into the brain. Cell Metab. 2014;19: 293–301.

19. Lazcano I, Cabral A, Uribe RM, Jaimes-Hoy L, Perello M, Joseph-Bravo P, et al. Fasting Enhances Pyroglutamyl Peptidase II Activity in Tanycytes of the Mediobasal Hypothalamus of Male Adult Rats. Endocrinology. 2015;156: 2713–2723.

20. Pak T, Yoo S, Miranda-Angulo AL, Wang H, Blackshaw S. Rax-CreERT2 knock-in mice: a tool for selective and conditional gene deletion in progenitor cells and radial glia of the retina and hypothalamus. PLoS One. 2014;9: e90381.

21. Imayoshi I, Sakamoto M, Ohtsuka T, Takao K, Miyakawa T, Yamaguchi M, et al. Roles of continuous neurogenesis in the structural and functional integrity of the adult forebrain. Nat Neurosci. 2008;11: 1153–1161.

22. Kokoeva MV, Yin H, Flier JS. Neurogenesis in the hypothalamus of adult mice: potential role in energy balance. Science. 2005;310: 679–683.

23. Pierce AA, Xu AW. De novo neurogenesis in adult hypothalamus as a compensatory mechanism to regulate energy balance. J Neurosci. 2010;30: 723–730.

24. Lee DA, Yoo S, Pak T, Salvatierra J, Velarde E, Aja S, et al. Dietary and sex-specific factors regulate hypothalamic neurogenesis in young adult mice. Front Neurosci. 2014;8: 157.

25. Campbell JN, Macosko EZ, Fenselau H, Pers TH, Lyubetskaya A, Tenen D, et al. A molecular census of arcuate hypothalamus and median eminence cell types. Nat Neurosci. 2017;20: 484–496.

26. Yoo S, Cha D, Kim DW, Hoang TV, Blackshaw S. Tanycyte-Independent Control of Hypothalamic Leptin Signaling. Front Neurosci. 2019;13: 240.

27. Song J, Christian KM, Ming G-L, Song H. Modification of hippocampal circuitry by adult neurogenesis. Dev Neurobiol. 2012;72: 1032–1043.

28. Miranda-Angulo AL, Byerly MS, Mesa J, Wang H, Blackshaw S. Rax regulates hypothalamic tanycyte differentiation and barrier function in mice. J Comp Neurol. 2014;522: 876–899.

29. Yasuo S, Yoshimura T, Ebihara S, Korf H-W. Temporal dynamics of type 2 deiodinase expression after melatonin injections in Syrian hamsters. Endocrinology. 2007;148: 4385–4392.

30. Sánchez E, Vargas MA, Singru PS, Pascual I, Romero F, Fekete C, et al. Tanycyte pyroglutamyl peptidase II contributes to regulation of the hypothalamic-pituitary-thyroid axis through glial-axonal associations in the median eminence. Endocrinology. 2009;150: 2283–2291.

31. de Seranno S, d’Anglemont de Tassigny X, Estrella C, Loyens A, Kasparov S, Leroy D, et al. Role of estradiol in the dynamic control of tanycyte plasticity mediated by vascular endothelial cells in the median eminence. Endocrinology. 2010;151: 1760–1772.

32. de Vries EM, Nagel S, Haenold R, Sundaram SM, Pfrieger FW, Fliers E, et al. The Role of Hypothalamic NF-κB Signaling in the Response of the HPT-Axis to Acute Inflammation in Female Mice. Endocrinology. 2016;157: 2947–2956.

33. Marsili A, Sanchez E, Singru P, Harney JW, Zavacki AM, Lechan RM, et al. Thyroxine-induced expression of pyroglutamyl peptidase II and inhibition of TSH release precedes suppression of TRH mRNA and requires type 2 deiodinase. J Endocrinol. 2011;211: 73–78.

34. Elizondo-Vega R, Cortés-Campos C, Barahona MJ, Carril C, Ordenes P, Salgado M, et al. Inhibition of hypothalamic MCT1 expression increases food intake and alters orexigenic and anorexigenic neuropeptide expression. Sci Rep. 2016;6: 33606.

35. Barahona MJ, Llanos P, Recabal A, Escobar-Acuña K, Elizondo-Vega R, Salgado M, et al. Glial hypothalamic inhibition of GLUT2 expression alters satiety, impacting eating behavior. Glia. 2018;66: 592–605.

36. Helfer G, Barrett P, Morgan PJ. A unifying hypothesis for control of body weight and reproduction in seasonally breeding mammals. J Neuroendocrinol. 2019;31: e12680.

37. Sáenz de Miera C. Maternal photoperiodic programming enlightens the internal regulation of thyroid-hormone deiodinases in tanycytes. J Neuroendocrinol. 2019;31: e12679.

38. Dardente H, Lomet D. Photoperiod and thyroid hormone regulate expression of l-dopachrome tautomerase (Dct), a melanocyte stem-cell marker, in tanycytes of the ovine hypothalamus. J Neuroendocrinol. 2018;30: e12640.

39. Kobayakawa K, Kobayakawa R, Matsumoto H, Oka Y, Imai T, Ikawa M, et al. Innate versus learned odour processing in the mouse olfactory bulb. Nature. 2007;450: 503–508.

40. Salvatierra J, Lee DA, Zibetti C, Duran-Moreno M, Yoo S, Newman EA, et al. The LIM homeodomain factor Lhx2 is required for hypothalamic tanycyte specification and differentiation. J Neurosci. 2014;34: 16809–16820.

41. Shimogori T, Lee DA, Miranda-Angulo A, Yang Y, Wang H, Jiang L, et al. A genomic atlas of mouse hypothalamic development. Nat Neurosci. 2010;13: 767–775.

42. Hübschle T, Thom E, Watson A, Roth J, Klaus S, Meyerhof W. Leptin-induced nuclear translocation of STAT3 immunoreactivity in hypothalamic nuclei involved in body weight regulation. J Neurosci. 2001;21: 2413–2424.

43. Vaisse C, Halaas JL, Horvath CM, Darnell JE Jr, Stoffel M, Friedman JM. Leptin activation of Stat3 in the hypothalamus of wild-type and ob/ob mice but not db/db mice. Nat Genet. 1996;14: 95–97.

