## Supplementary material for "Ablation of tanycytes of the arcuate nucleus and median eminence increases visceral adiposity and decreases insulin sensitivity in male mice": Supplemetal Figures S1-S7

**Supplementary figure 1. Hypothalamic neuronal distribution and size of nuclei within reach of tanycytes. (A)** *Npy*, *Cart*, *Gal* and *Sst* mRNA expression in tubular hypothalamic region of wildtype and tanycyte-ablated mice. **(B)** Quantification of the size of hypothalamic nuclei located adjacent to tanycytic layer. **(C)** Representative images of anti-Neurofilament M subunit immunostaining in the ME. **(D)** *Oxt*, *Trh* and *Avp* mRNA expression in anterior hypothalamic regions where neurons project to or pass through the ME. DMH, dorsomedial nucleus; VMH, ventromedial nucleus; ArcN, arcuate nucleus; ME, median eminence; 3V, third ventricle. Scale bar: 200 µm (A,D), 100 µm (C).

**Supplementary figure 2. Effect of tanycyte-ablation on food intake, metabolic hormone level and glucose/insulin sensitivity in female mice. (A, B)** *Ad libitum* (*Ad lib*) or 18h fasting-induced (re-feed) food intake at 5 weeks (A) and 8 weeks (B) after starting tamoxifen diet. **(C)** Serum leptin and insulin level measured by Luminex assay using metabolic bead panel. **(D)** Glucose tolerance test in long-term tamoxifen fed mice. **(E)** Insulin tolerance test in long-term tamoxifen fed mice. $^*p<0.02$.

**Supplementary figure 3. Serum hormone levels in long-term tamoxifen fed male mice. (A-F)** Effects of ArcN-ME tanycyte-ablation in serum concentration of metabolic hormones, PP (pancreatic polypeptide), PYY (peptide YY), C-Peptide, GIP (gastric inhibitory polypeptide), TNF-α (tumor necrosis factor alpha), Testosterone. **(G-M)** Effects of ArcN-ME tanycyte-ablation in serum concentration of pituitary hormones, ACTH (adrenocorticotropic hormone), FSH (follicle stimulating hormone), LH (luteinizing hormone), TSH (thyroid-stimulating hormone), GH (growth hormone), PRL (prolactin), Oxytocin. **(N-Q)** Effects of ArcN-ME tanycyte-ablation in serum concentration of neuropeptides, α-MSH (alpha melanocyte stimulating hormone), Neurotensin, Substance P, β-Endorphin. $^*p<0.05$, $^{**}p<0.01$.

**Supplementary figure 4. Serum hormone levels in short-term tamoxifen fed male mice. (A-F)** Effects of ArcN-ME tanycyte-ablation in serum concentration of metabolic hormones, PP (pancreatic polypeptide), PYY (peptide YY), C-Peptide, GIP (gastric inhibitory polypeptide), TNF-α (tumor necrosis factor alpha), Testosterone. **(G-M)** Effects of ArcN-ME tanycyte-ablation in serum concentration of pituitary hormones, ACTH (adrenocorticotropic hormone), FSH (follicle stimulating hormone), LH (luteinizing hormone), TSH (thyroid-stimulating hormone), GH (growth hormone), PRL (prolactin), Oxytocin. **(N-Q)** Effects of ArcN-ME tanycyte-ablation in serum concentration of neuropeptides, α-MSH (alpha melanocyte stimulating hormone), Neurotensin, Substance P, β-Endorphin. $^*p<0.04$.

**Supplementary figure 5. Serum hormone levels in long-term tamoxifen fed female mice. (A-E)** Effects of ArcN-ME tanycyte-ablation in serum concentration of metabolic hormones, PP (pancreatic polypeptide), PYY (peptide YY), C-Peptide, GIP (gastric inhibitory polypeptide), TNF-α (tumor necrosis factor alpha), Testosterone. **(F-L)** Effects of ArcN-ME tanycyte-ablation in serum concentration of pituitary hormones, ACTH (adrenocorticotropic hormone), FSH (follicle stimulating hormone), LH (luteinizing hormone), TSH (thyroid-stimulating hormone), GH (growth hormone), PRL (prolactin), Oxytocin. **(M-P)** Effects of ArcN-ME tanycyte-ablation in serum concentration of neuropeptides, α-MSH (alpha melanocyte stimulating hormone), Neurotensin, Substance P, β-Endorphin. *$p<0.003$.

**Supplementary figure 6. Effect of tanycyte-ablation on serum level of TSH in animals exposed to acute cold. (A)** TSH serum concentration before and after a cold challenge for 2 hours at 4°C in wildtype (WT) and tanycyte-ablated mice (Tny-DTA). **(B)** Relative change in TSH serum level calculated from the result in (A). *$p<0.04$, **$p<0.001$.

**Supplementary figure 7. Indirect calorimetry analysis in tanycte-ablated mice.** **(A)** Experimental schedule for indirect calorimetry after dietary tamoxifen administration. **(B)** Body weights of animals subjected to indirect calorimetry. **(C)** Food intake during metabolic profiling. **(D-F)**. Respiratory exchange ratio (RER, $VCO_2/VO_2$, D), oxygen consumption ($VO_2$, E), and energy expenditure (F) measured in wildtype (WT) and tanycyte-ablated (Tny-DTA) mice. *, significant difference ($p<0.05$) between groups continued more than 30min.

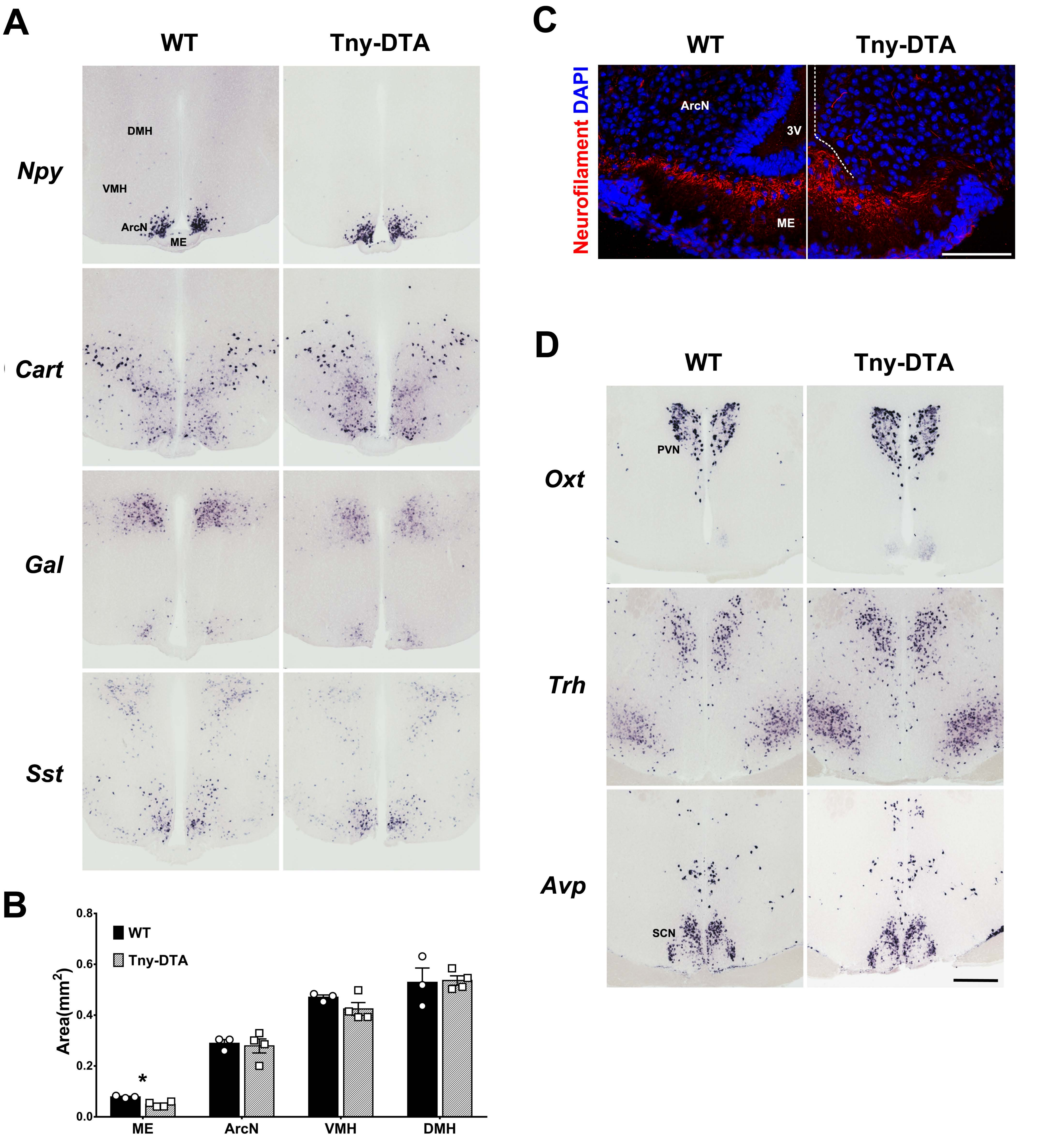


Yoo, et al. Figure S1

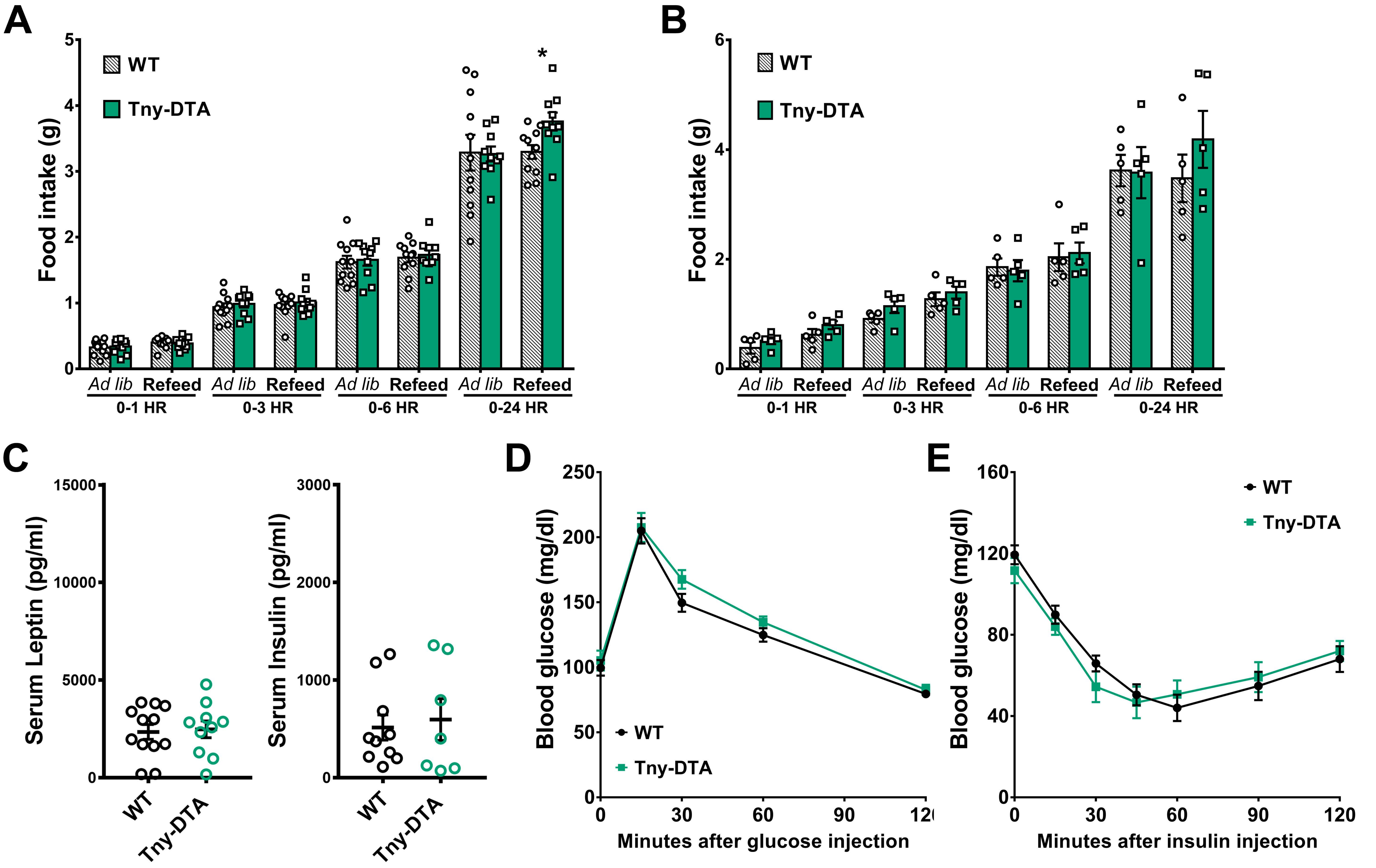


Yoo, et al. Figure S2

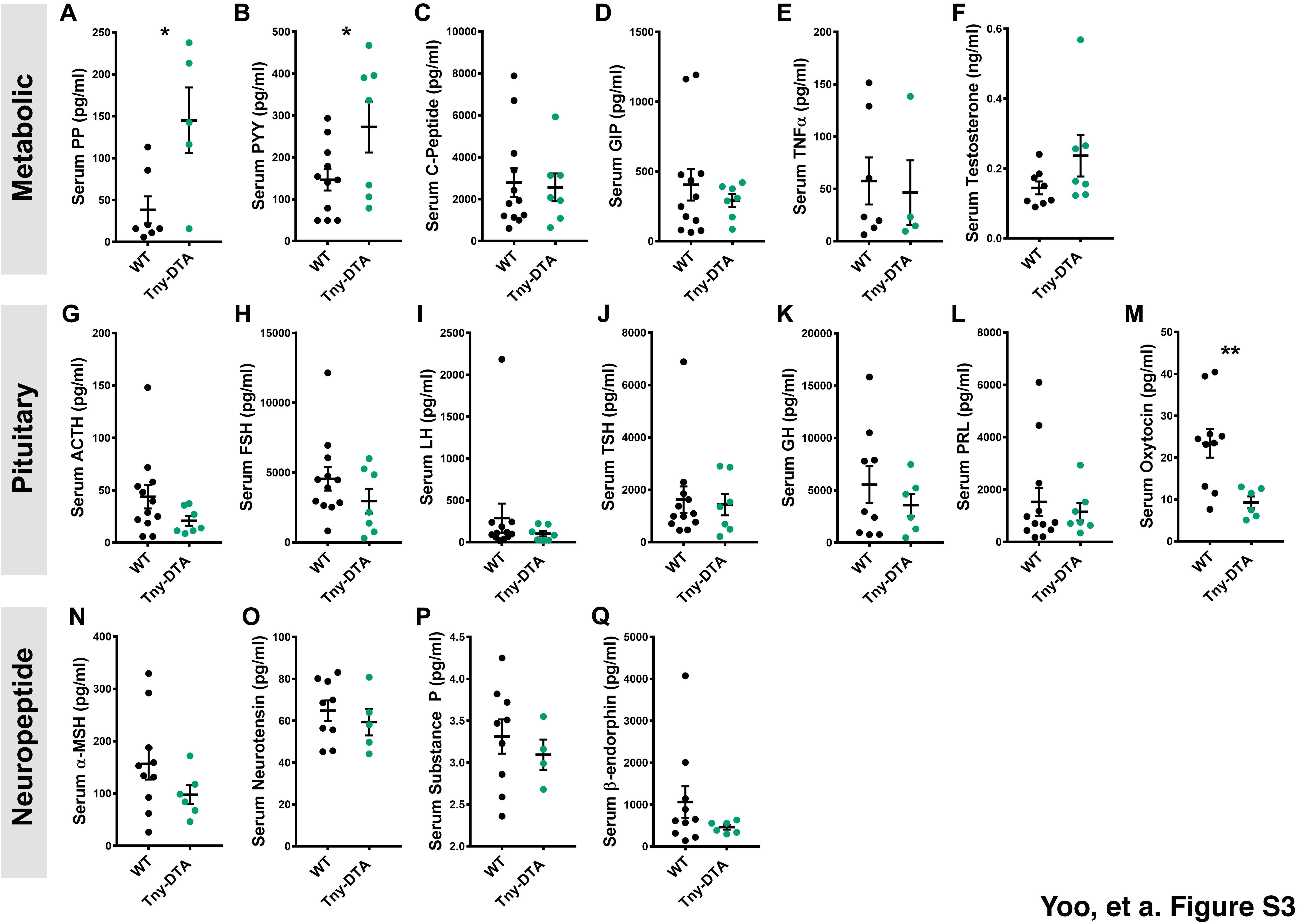


Yoo, et a. Figure S3

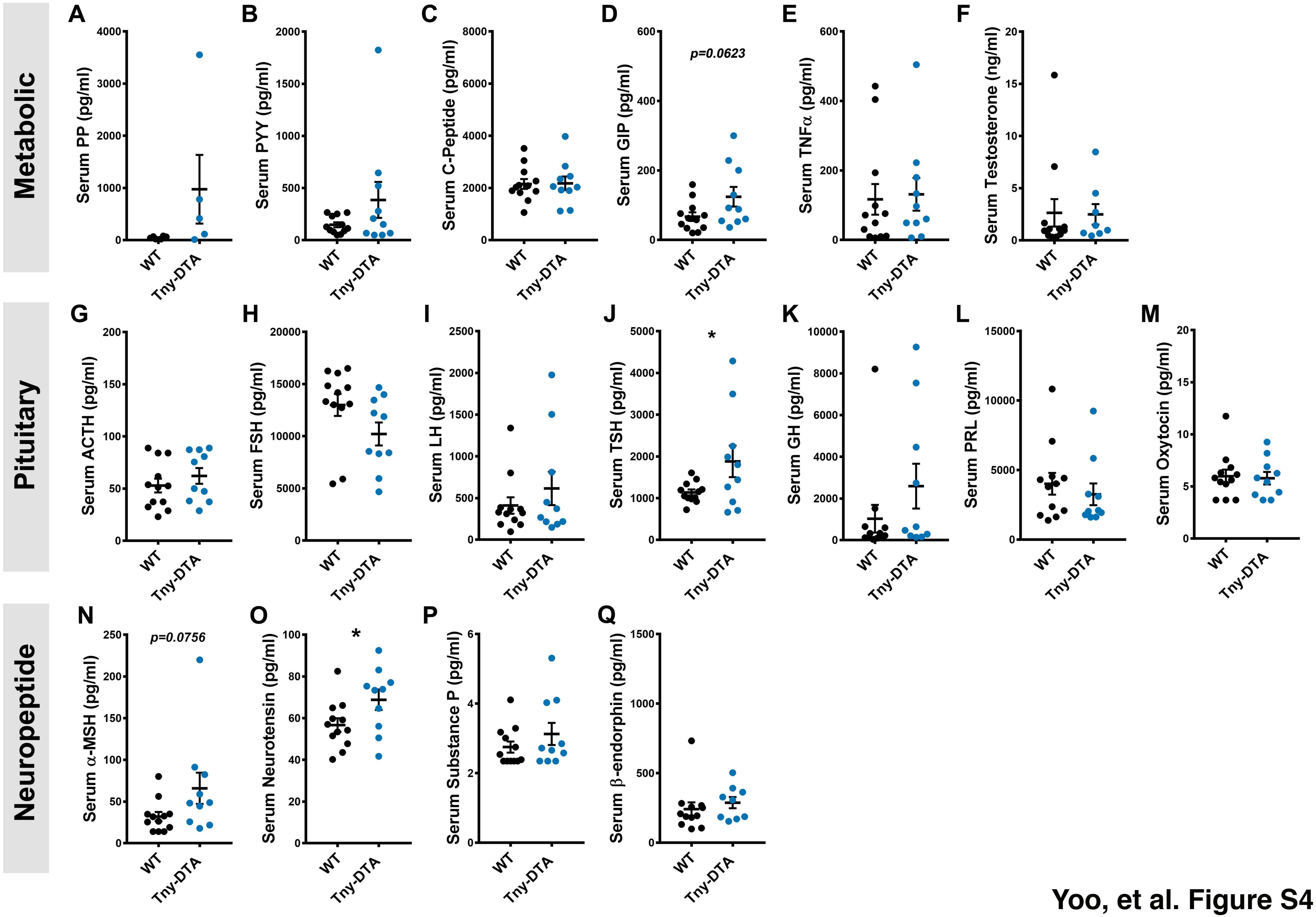


Yoo, et al. Figure S4

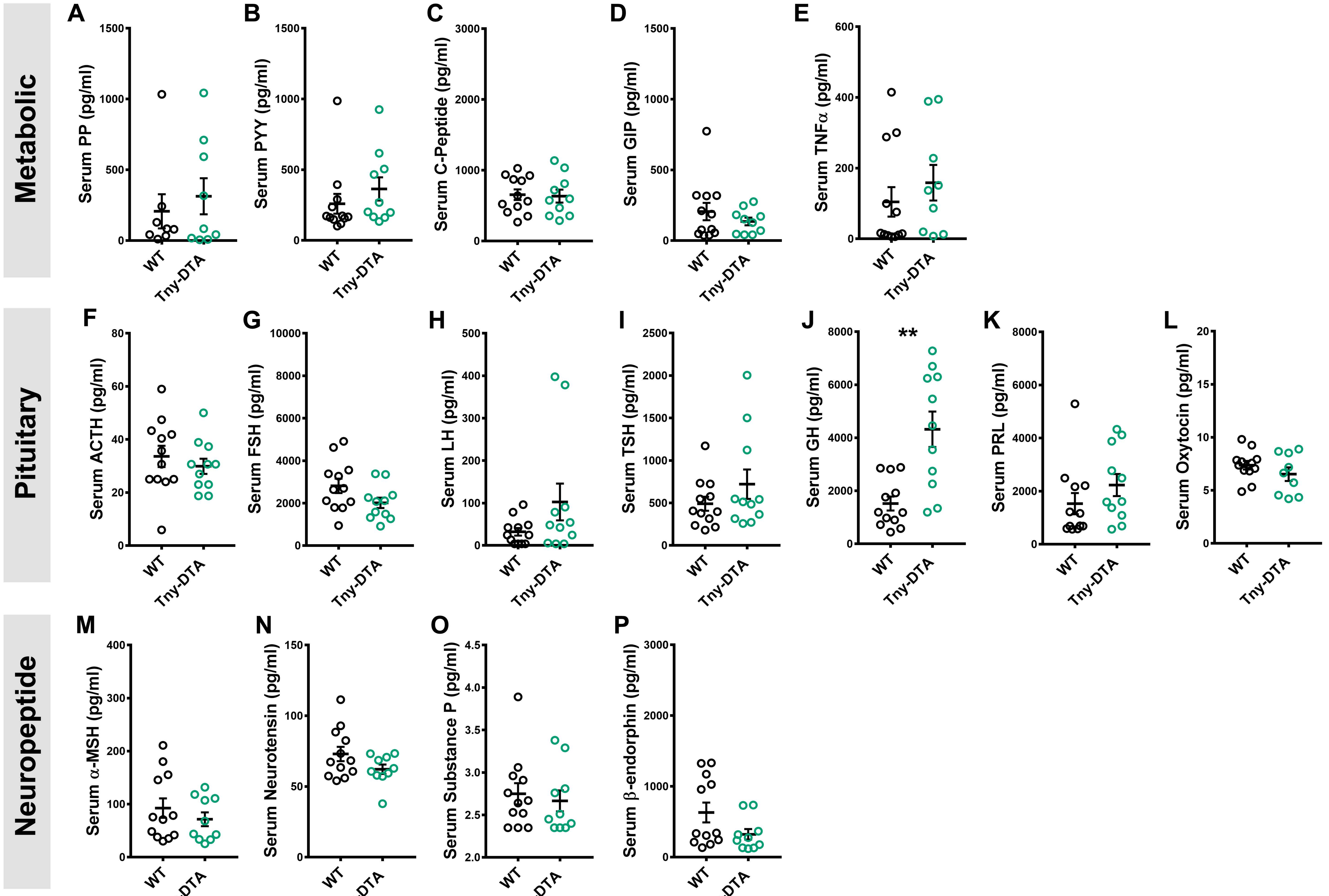


Yoo, et al. Figure S5

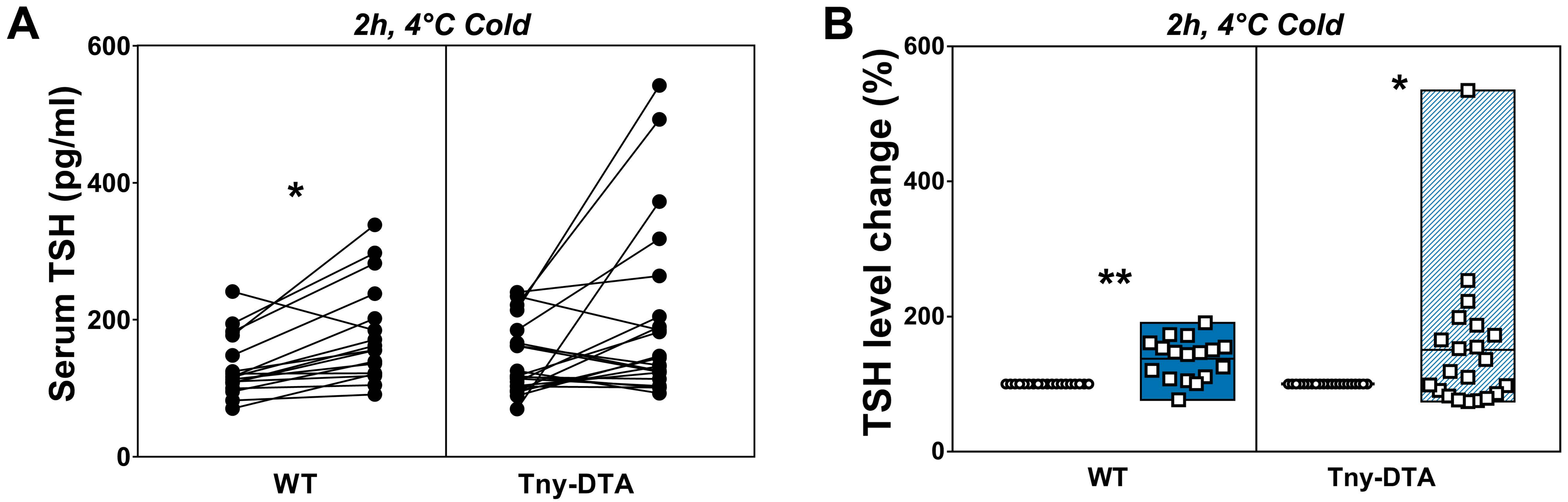


Yoo, et al. Figure S6

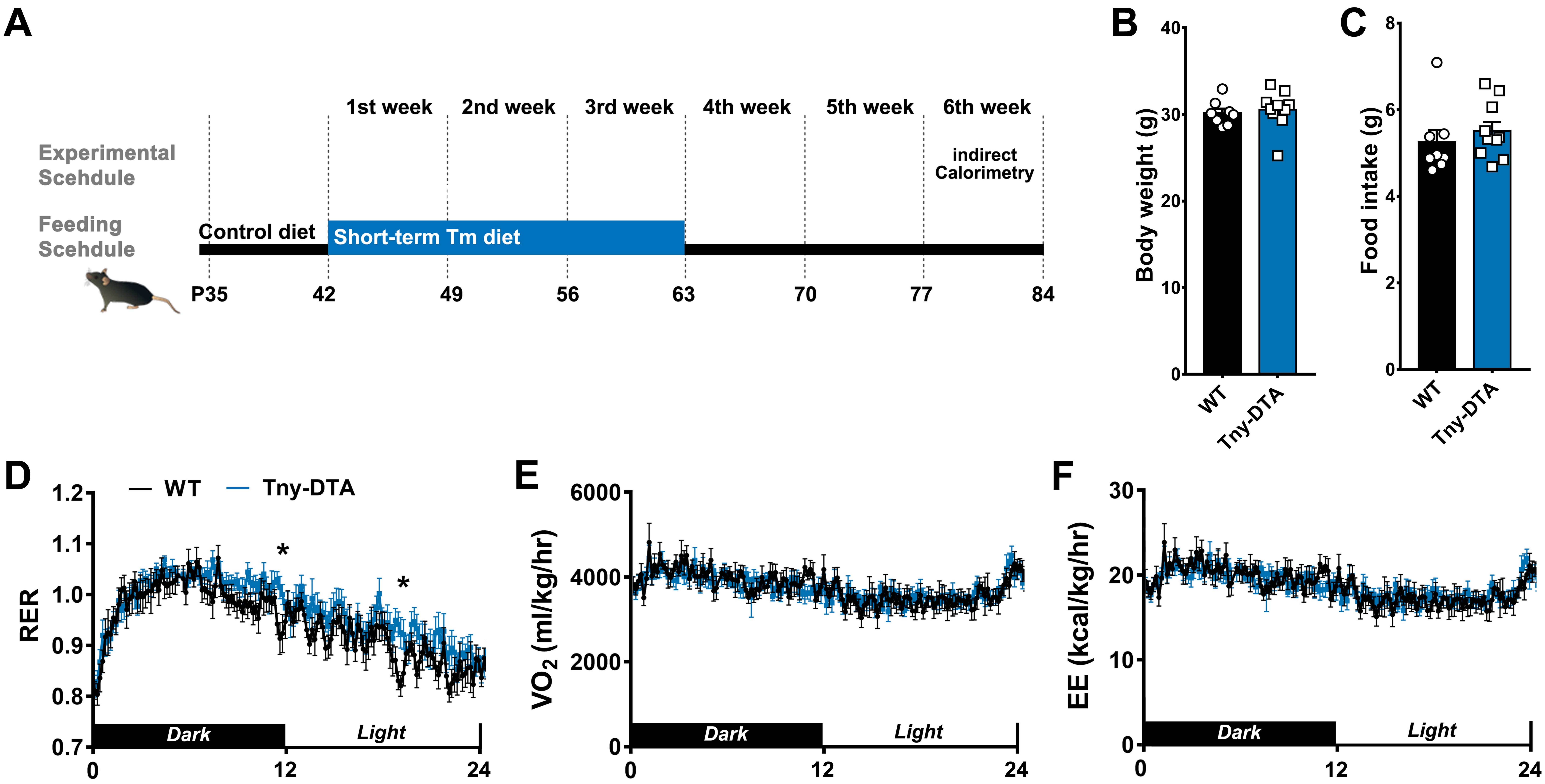


Yoo, et al. Figure S7
